# The allotetraploid *Nicotiana tabacum* genome and GenBank genomics highlight the genomic features, genetic diversity and regulation of morphological, metabolic and disease-resistance traits

**DOI:** 10.1101/2023.02.21.529366

**Authors:** Yanjun Zan, Shuai Chen, Min Ren, Guoxiang Liu, Yutong Liu, Huan Si, Zhengwen Liu, Dan Liu, Xingwei Zhang, Ying Tong, Yuan Li, Caihong Jiang, Liuying Wen, Zhiliang Xiao, Yangyang Sun, Ruimei Geng, Quanfu Feng, Yuanying Wang, Guoyou Ye, Lingzhao Fang, Yong Chen, Lirui Cheng, Aiguo Yang

## Abstract

*Nicotiana tabacum* is a model organism in plant molecular and pathogenic research and has significant potential in the production of biofuels and active pharmaceutical compounds in synthetic biology. Because of the large allotetraploid genome of tobacco, its genomic features, genetic diversity and genetic regulation of many complex traits remain unknown. In this study, we present a nearly complete chromosome-scale assembly of *N. tabacum* and provide evidence that homoeologous exchange between subgenomes and epigenetic remodelling are likely mechanisms of genome stabilization and subgenome coordination following polyploidization. By leveraging GenBank-scale sequencing and phenotyping data from 5196 lines, geography at the continent scale, rather than types assigned on the basis of curing crop practices, was found to be the most important correlate of genetic structure. Using 178 marker□trait associations detected in genome-wide association analysis, a reference genotype-to-phenotype map was built for 39 morphological, developmental, and disease-resistance traits. A novel gene, auxin response factor 9 (*Arf9*), associated with wider leaves after being knocked out, was fine-mapped to a single nucleotide polymorphism (SNP). This point mutation alters the translated amino acid from Ala_203_ to Pro_203_, likely preventing homodimer formation during DNA binding. Our analysis also revealed signatures of positive and polygenic selection for multiple traits during the process of selective breeding. Overall, this study demonstrated the power of leveraging GenBank genomics to gain insights into the genomic features, genetic diversity, and regulation of complex traits in *N. tabacum*, laying a foundation for future research on plant functional genomics, crop breeding, and the production of biopharmaceuticals and biofuels.

## Introduction

Nicotiana species have served as models for a series of cytogenetic and molecular genetic breakthroughs, including the discovery of protoplast fusion^1^, the introduction of foreign genetic material through interspecies hybridization^2,3^, T-DNA mediated plant transformation^4^, virus-induced genetic transformation^5^, the first edible vaccine^6,7^, and plant pathogen studies^8^. The rich diversity in plant architecture, leaf morphology, flowering time and metabolic traits, ease of genetic manipulation, and fairly high number of curated pathways in *Nicotiana tabacum* highlight its central importance in plant development research and its significant potential in the field of synthetic biology^6,7^.

*Nicotiana tabacum* (2n=4x=48, ~4.30 Gb) is an interspecific hybrid derived from two progenitors, *Nicotiana sylvestris* (2n=2x=24, 2.59 Gb) and *Nicotiana tomentosiformis* (2n=2x=24, 2.22 Gb) approximately 0.2 Mya ago^9–11^. Several attempts have been made to assemble its large allotetraploid genome; however, the latest assembly included only 81% of the genomic sequences and was very fragmented^9,11,12^. As a consequence of the large and repetitive polyploid genome, obtaining a chromosome-level assembly constitutes a significant challenge, as there are difficulties not only in bridging highly repetitive repeat regions but also in computationally disentangling homologous sequences with high similarity from two progenitor species^9,11,12^. In the absence of a high-quality genome assembly and with nearly all genetic studies being performed in the private sector, the genomic features, global genetic diversity and genetic basis of many important traits in this model plant remain poorly understood. These factors highlight the necessity for a complete genome assembly and GenBank-scale holistic investigation of the genetic diversity and regulation of complex traits that are highly relevant for future plant functional genomics, crop improvement, and plant synthetic biology research.

In this study, we present the first complete chromosome-level assembly of the allotetraploid *Nicotiana tabacum* genome and genetic and phenotypic data for an entire *Nicotiana tabacum* GenBank with 5196 germplasms hosted at the Chinese Academy of Agriculture Sciences. The complete genome assembly demonstrated that chromosome rearrangement between subgenomes and epigenetic remodeling are likely mechanisms in genome stabilization and subgenome coordination following polyploidization. By utilizing this valuable resource, insights were gained into the genomic features, genetic diversity and genetic regulation of plant architecture, morphological, metabolic, and disease-resistance traits in this model species. A novel gene, auxin response factor 9 (*Arf9*), resulting in wider leaves after being knocked out, was fine-mapped to a single nucleotide polymorphism (SNP) altering the translated amino acid from Ala_203_ to Pro_203_, which in turn prevented the formation of a homodimer during DNA binding. Results from our study, together with the released data and seeds, will likely accelerate aspects of tobacco research that were previously hindered by the complexity of the polyploid genome and a lack of publicity for research driven by the private sector, with benefits in plant functional genomics, crop improvement research, and the production of biopharmaceutical compounds and biofuel.

## Results

### Chromosome-scale assembly and genomic features of the allotetraploid Nicotiana tabacum genome

Based on k-mer analysis with 52X Illumina short reads and flow cytometry analysis, the *N. tabacum* genome was estimated to be 4.29-4.30 Gb, representing a downsizing of approximately 10% from the sum of the two parental genomes (Fig 1A, Supplementary Fig S1). To overcome the challenge of assembling a large polyploid genome, we generated 52X Illumina short reads, 123.35 X PacBio Sequel II reads, 124.93X 10X genomic linked reads, 120.12 X Hi-C reads and 192.27 X Bionano reads from *N. tabacum L*. var *ZY300* and adopted a hybrid assembly approach (Materials and methods). The final assembly included 4.17 Gb (Fig 1B) of sequences with contig N50 at 27.17 Mb, and 96.98% of the sequences were anchored to 24 pseudochromosomes (Supplementary Table 1, Supplementary Fig S2). This represented a 10-15 percent increase in the assembled genome size and a 10-fold increase in contig N50 length compared to those of the latest assembly (Supplementary Table 1) ^9,11,12^. The continuity and completeness of the assembled genome were supported by a high mapping rate (99.72%) and high genome coverage (99.47%) of Illumina short reads, 99.2% single-copy orthologues from the Benchmarking Universal Single-Copy Orthologues (BUSCO) analysis and a high long terminal repeat (LTR) assembly index (LAI) score (LAI=9.74).

**Fig 1.**
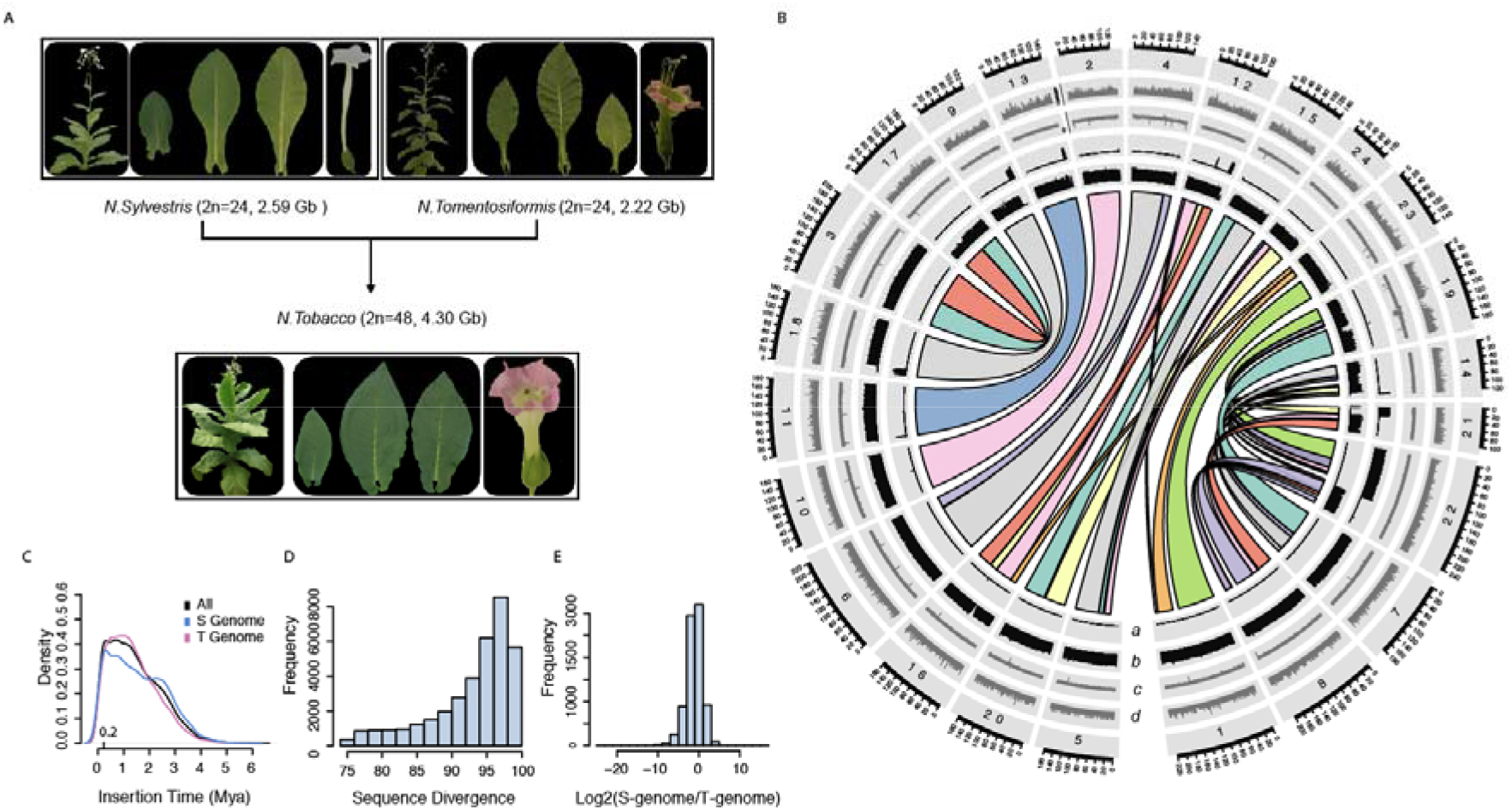
High-quality assembly and features of the *N. tabacum* genome. **A)** Phenotype diversity and genome size variation among two diploid progenitors, *N. sylvestris* and *N. tomentosiformis*, and one of the allotetraploid *N. tabacum* samples. **B)** Genomic features of *N. tabacum*. Tracks from innermost to outermost are sequence coverage obtained by mapping Illumina short reads from *N. tomentosiformis* (a) and *N. sylvestris* (b), gene density (c), and transposable element (TE) density (d). Synteny blocks between the two subgenomes are represented by coloured linkers across the centre of the plot. **C)** Estimated insertion time for TEs from the two subgenomes. Histogram of sequence divergence **D)** and expression divergence **E)** of 35740 protein-coding homologous gene pairs from the two subgenomes.

The complete assembly of a complex allotetraploid presents the opportunity to study subgenome evolution and coordination following polyploidization. In total, 57.14% and 42.85% of the genome was partitioned to *N. sylvestris* (S) and *N. tomentosiformis* (T), respectively, with 12 chromosome rearrange events pinpointed with approximately 1 kb resolution (Supplementary Table 2). This suggested that chromosomal rearrangements produced significant changes in genome structure, which likely stabilized chromosome pairing after polyploidization. To evaluate the mechanisms contributing to genome downsizing and subgenome coordination, we annotated the transposable elements (TEs) and homologous gene pairs and quantified their sequence and expression divergence. Based on a de novo-constructed species-specific TE library, 82.75% of the genome was masked. Estimation of the TE insertion time showed that approximately 7.56% of the TEs were inserted after merging of the two subgenomes 0.2 Mya ago (Fig 1C), suggesting that polyploidization did not stimulate extensive reactivation of TEs. The overall sequence similarity of 35740 homologous gene pairs between subgenomes was high (median = 92%, Fig 1D), and expression divergence was low (median fold change estimated from 400 RNA-seq data points of 0.98, Fig 1E, Supplementary Table 3). However, over a thousand homologous genes with high sequence similarity displayed considerable expression divergence (median fold change = 3.92, Fig 1E, Supplementary Table 3, Supplementary Fig S3), indicating a likely role of epigenetic remodelling in subgenome gene expression coordination for a small proportion of homologous genes. A detailed summary of the TE annotation and gene annotation statistics is available in Supplementary Tables 4-5.

### GenBank genomics highlights the diversity of a global collection of 5196 tobacco lines

To understand the global distribution of genetic variation and genetic structure, we genotyped an entire GenBank collection hosted at the Chinese Academy of Agriculture Sciences using a genotype-by-sequencing approach (Materials and methods). This is one of the world’s largest collections of tobacco germplasm resources, with 2582 sun-cured tobacco, 2152 flue-cured tobacco, 223 burley tobacco, 126 cigar tobacco and 113 oriental tobacco germplasms, amounting to 5196 germplasms in total (Supplementary Table 6). These five types of tobacco were named after their roles in agronomical practices (Supplementary Table 7) and characterized by distinctive characteristics in plant architecture, leaf morphology, flowering time and metabolic traits (Fig 2A, Supplementary Fig S4).

**Fig 2.**
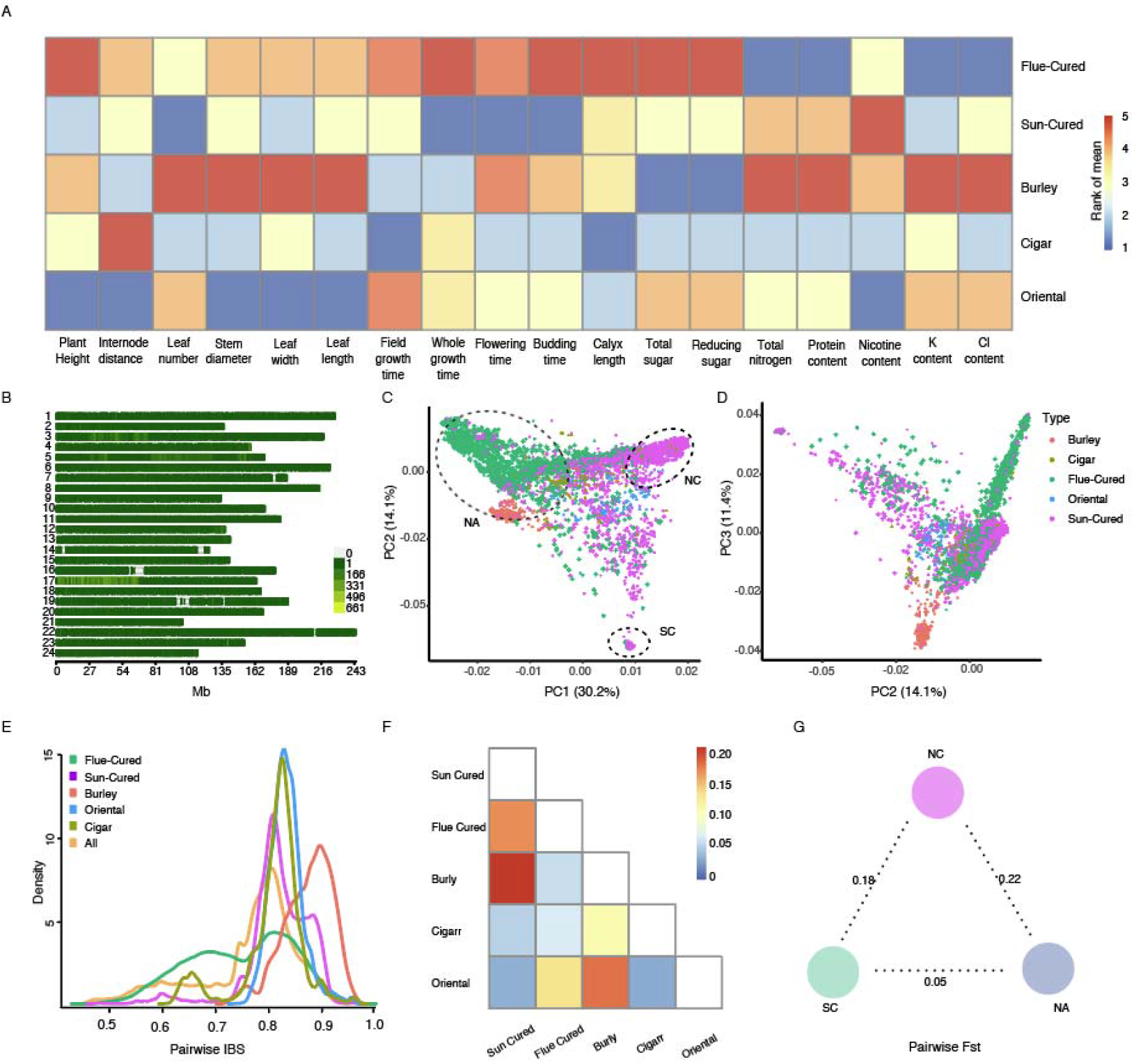
The genetic and phenotypic diversity of a global collection of 5,196 tobacco germplasm resources. **A).** A heatmap of mean phenotype rank for 20 traits (x-axis) among five types of tobacco named on the basis of their role in agronomical practices. **B).** Genomic distribution of 95,308 SNPs in the *N. tabacum* genome called with the genotype-by-sequencing approach. **C), D)** Genetic structure of 5196 *N. tabacum* lines inferred from principal component analysis. Samples are coloured according to the type assigned on the basis of agronomical practices. **E).** Distribution of pairwise IBS values in different tobacco types. Fixation index (Fst) between different tobacco types **F).** Groups defined on the basis of genetic structure **G).**

Using 95,308 SNPs covering 98% of the genomic bins at a 1 Mb resolution (Fig 2B), we found that geography at the continent scale was the most important correlate of genetic structure and that genetic structure did not always match the conventional types defined on the basis of agronomical practices. For example, the first principle component (PC1) separated flue-cured tobacco from sun-cured tobacco to a large extent. However, the majority of samples located on the left side of Fig 2C were of flue-cured tobacco originating from North America (NA) or derived materials belonging to the NA lineage, while plants located on the opposite side were sun-cured tobacco landraces collected in China (Fig 2C/2D). In addition, a latitudinal cline was observed along the second principal component (PC2). Sun-cured tobacco landraces from northern China (NC) with a significantly shorter life cycle, colder growth temperature and longer day length were located in the upper right corner (Cluster NC, Fig 2C), while plants sampled in southern China (SC) with opposite climate signatures were clustered at the bottom of Fig 2C. According to historical records, burley tobacco originated from a single mutated variety, known as White Burley, characterized by a chlorophyll deficiency phenotype controlled by double homozygous recessive alleles at the Yellow burley 1 (*YB1*) and Yellow burley 2 (*YB2*) loci^13^. Consistent with this, burley tobacco samples were clustered together near the NA group and showed the highest pairwise isolation by distance (IBS) value (median=0.88, Fig 2E). Cigar and oriental tobacco samples were scattered among sun-cured and flue-cured tobacco samples, indicating a likely admixed breeding history (Fig 2C/D).

Pairwise IBS values for the five tobacco types ranged from 0.75-0.88, suggesting that the gene pool used to develop each type of tobacco was very narrow. However, the distribution of IBS values for flue-cured, sun-cured and cigar tobacco was multimodal, indicating the cooccurrence of divergent gene pools likely due to introgression from foreign materials. Despite the marked phenotypic differentiation in plant morphology, flowering and metabolic traits (Fig 2A/Supplementary Fig S4), genome-wide differentiation among the five types and three genetic groups was low, with pairwise F_st_ values ranging from 0.04 to 0.23 (Fig 2F/G) and no selective sweeps detected (data not shown here). This is most likely due to the short and special selective breeding history of less than a few hundred years aiming to balance quality among many leaf characteristics.

### A comprehensive catalogue of the allelic variations underlying variation of plant morphological, physiological, metabolic and disease-resistance traits

With the availability of genome-wide markers and historical phenotypic records for over 5000 lines, we used the genome-wide association study (GWAS) approach to generate a comprehensive catalogue of the allelic variation underlying differences in 43 traits (Fig 2A/Fig 3A) with intermediate to high heritability (*h^2^* = 0.15-0.82, Supplementary Tables 8-9). A total of 178 significant marker□trait associations were identified for 39 traits (Fig 3B, Supplementary Fig S5), and several high-potential associations were detected and explored further. P values and additive effects for the quantitative trait loci (QTLs) are summarized in Supplementary Table 10. Although linkage disequilibrium (LD) is too extensive to directly pinpoint the candidate genes underlying each association peak, we found a few genes previously identified as QTLs^12,14–18^ or annotated as potential candidates near eight peaks (dashed lines in Fig 3B). In addition, we conducted a review of the literature^12,14–18^ to determine the relative position of previously reported QTLs (Supplementary Table 11) in the ZY300 genome assembly and discovered that 95% of the QTLs detected in our study were novel. Altogether, these marker□trait associations constitute a comprehensive genotype-to-phenotype map (Fig 3D, Supplementary Table 12), providing a roadmap for future genetic studies in this model plant species.

**Fig 3.**
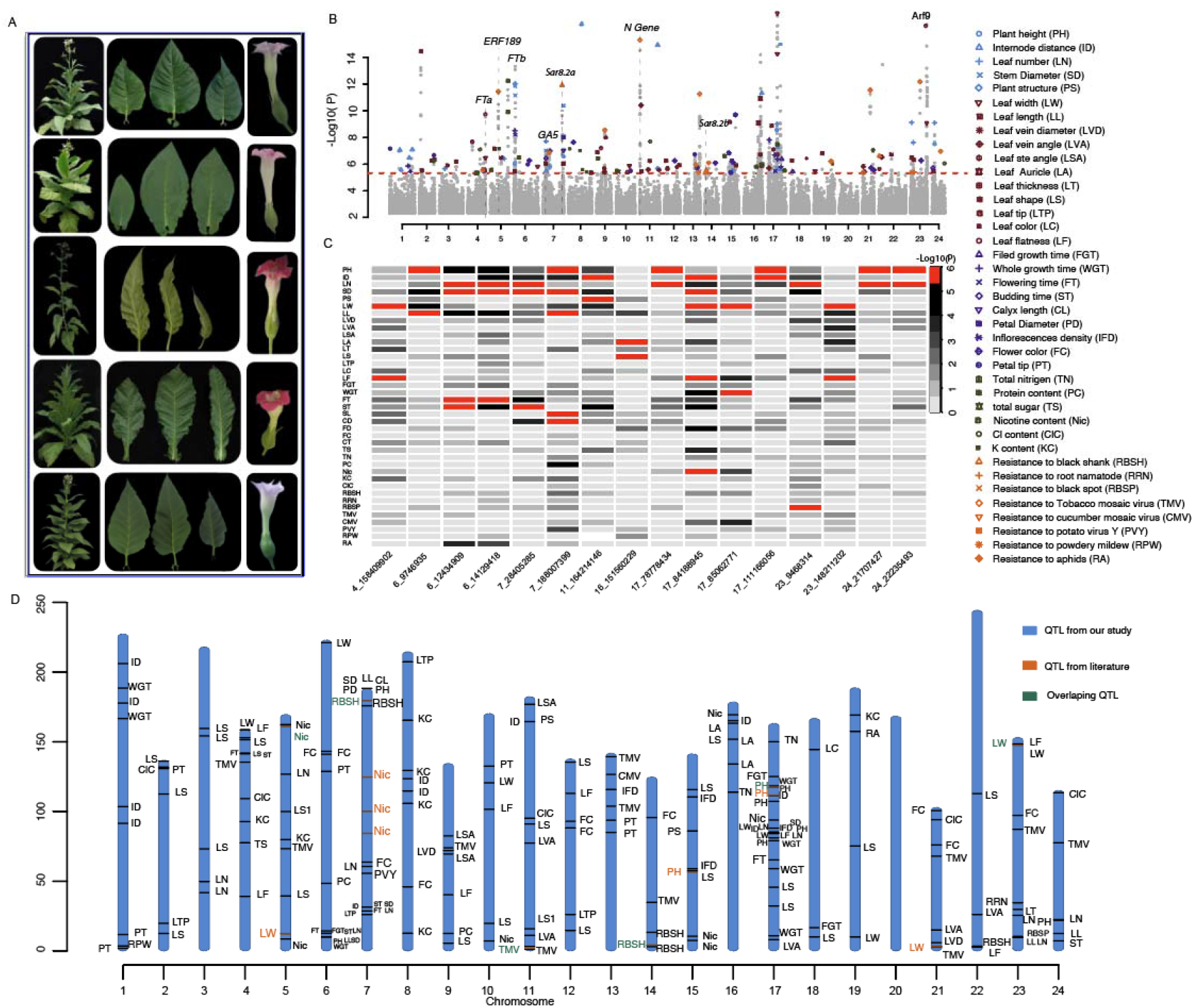
Association results for 43 plant morphological, physiological, metabolic and disease-resistance traits. **A).** Phenotype diversity of plant architecture, leaf morphology and flowering morphology and colour among five selected plants. The vertical dashed grey lines highlight the genomic position of potential candidate genes. **B).** Manhattan plots of GWAS results from 43 GWAS scans. The red horizontal dashed lines indicate the Bonferroni-corrected genome-wide significance thresholds. The vertical dashed grey lines highlight the genomic position of 16 SNPs associated with more than two traits. **C).** A heatmap illustrating the p values of 16 SNPs detected for more than two traits. Each cell represents the −log10 (p value) of a particular SNP (x-axis) associated with a specific trait (y-axis on the right). **D).** The reference genotype-to-phenotype map includes marker□trait associations detected in our study (orange), previous reports (sky blue), and both (dark green). Previously reported QTLs spanning more than 10 Mb and QTLs without probe sequences are not included here.

To make these resources accessible to a wide range of researchers, we selected 310 accessions to cover the majority of the genetic and phenotypic diversity, and have made their seeds available for ordering (See for detailed information). Our intention is for this collection to remain actively curated as ever more genomic data are produced and a wide range of phenotypic data are generated (not only by us, but also by the community— see for detailed information on those accessions and on how to contribute).

We detected 32 QTLs (Supplementary Tables S10/S12) for four plant architecture traits, namely, plant height (PH), internode distance (ID), leaf number (LN) and stem diameter (SD), that were moderately correlated (Fig 3B, Supplementary Fig S6). Overall, taller plants tended to have more leaves, wider stems and larger ID values (Fig 4A, Supplementary Fig S4), indicating a shared molecular basis for plant architecture development. Consistent with this finding, four QTLs were associated with variation in two or more plant architecture traits (Fig 3B/3C). For example, the QTL (Chr7: 28 405 285 bp) located at the proximal end of chromosome 7 was associated with variation in PH, LN and SD (Fig 4B). Allele G at this locus showed a dominance effect over allele A (Fig 4D), whereby the GG genotype at this locus increased ID, LN and SD by 0.22 ± 0.04 cm (P= 6.32 x 10^-6^), 0.79 ± 0.15 (P= 4.02 x 10^-8^) and 0.29 ± 0.05 cm (P= 8.60 x 10^-11^), respectively (Fig 4D/E/F). Although the LD in this region extends for several megabases, we discovered a potential candidate, NtZY07G00865, homologous to *Arabidopsis thaliana* gibberellin 20-oxidase (AT4G25420, GA5), which regulates plant developmental processes^19^, located within the association peaks.

**Fig 4.**
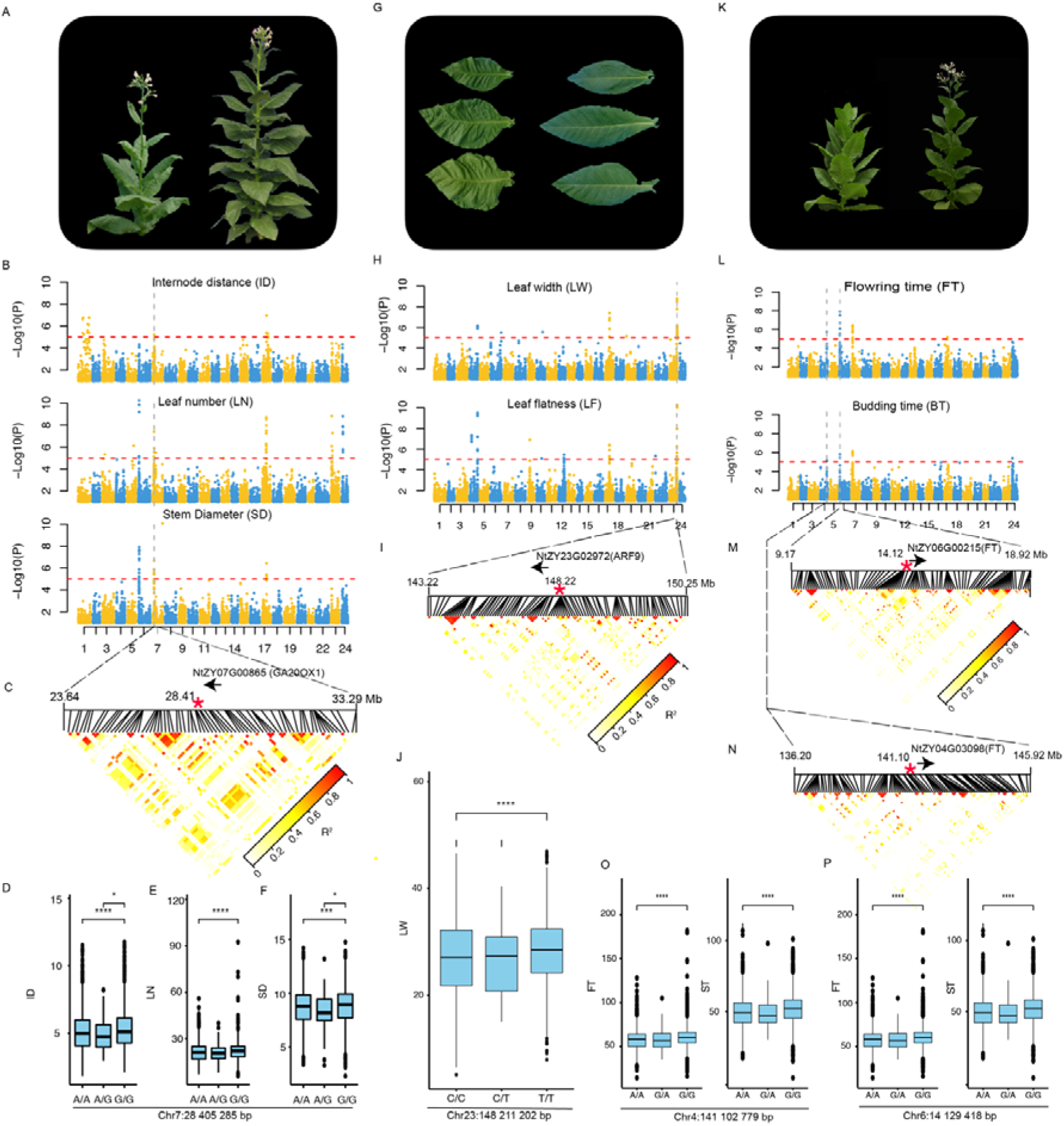
GWASs identify potential candidate genes for plant architecture, leaf morphology and flowering time variation. Graphical illustration of the variation in plant architecture **A),** leaf morphology **G),** and flowering time **K)** traits in *Nicotiana tobacco*. Association results for internode distance, stem diameter and leaf number **B)**, leaf width, leaf flatness **H),** flowering time and bolting time **L). C).** The LD heatmap and literature review pinpointed NtZY07G00865, a homologue of *A. thaliana* AT4G25420 (*GA5*) involved in the later steps of the gibberellin biosynthetic pathway, as a potential candidate underlying the association peak at chromosome 7: 28 405 285 bp. **I).** The LD heatmap and literature review pinpointed NtZY23G02972, a homologue of *A. thaliana* auxin response factor 9 (*Arf9*), which regulates cell division activity, as a potential candidate underlying the association peak at chromosome 23: 148 211 202 bp. **M), N),** The LD heatmap and literature review pinpointed NtZY04G03098 and NtZY06G00215, both homologous to *A. thaliana* flowering time locus T involved in flowering time regulation, as potential candidates underlying the association peak at Chr6:14 129 418 bp and Chr4:141 102 779 bp, respectively. **D), E), F), J), O), P)** Genotype-to-phenotype map for the QTLs associated with the corresponding traits.

Strong agronomical interests in leaf usage resulted in an excess amount of diversity in leaf morphology (Fig 3A, Fig 4G) traits. Here, we detected 65 QTLs for 12 leaf morphology traits (Supplementary Table S8), including leaf width (LW), leaf length (LL), vein diameter (VD), leaf stem angle (LSA), leaf vein angle (LVA), leaf auricle (LA), leaf thickness (LT), leaf shape (LS), leaf tip (LTP), leaf colour (LC), leaf serration (LSR), and leaf flatness (LF). Overall, the correlations between these traits were relatively low (median Spearman correlation = 0.05, Fig S3). However, 3 QTLs were simultaneously associated with LW and LF (Fig 3B/Fig 4H). One of these QTLs (Chr23: 148 211 202 bp) is located at the end of chromosome 23. Allele T at this locus increased LW by 1.11 ± 0.18 (P= 1.66 x 10^-9^, Fig 4J) and increased LF by 0.2 ± 0.02 units (P = 1.58 x 10^-18^). Similar to that for the peak described in the previous section, the LD in this region was too extensive to directly pinpoint candidates; however, we found NtZY23G02972, a homologue of *A. thaliana* auxin response factor 9 (*Arf9*), which regulates cell division activity, to be a potential candidate for further validation (Fig 4I).

Regulation of flowering time variation has attracted broad interest due to its importance in plant development, ecological adaptation and breeding applications. Here, four QTLs were detected for flowering time (FT) and budding time (BT) variation (Fig 4 K-L). Among these, two QTLs located at the end of chromosome 4 (Chr4:141 102 779 bp) and proximal end of chromosome 6 (Chr6:14 129 418 bp) attracted our attention (Fig 4L). Both QTLs included a candidate gene homologous to *A. thaliana* flowering time locus T (*FT*), and these two chromosome regions were synteny blocks from the S and T subgenomes (Fig 4 M/N, Fig 1B). This suggests that the function of both homologous genes was retained after merging of the two subgenomes.

In addition to the association peaks described above, we detected additional QTLs associated with leaf metabolites and resistance to several virus-/bacterium-/fungus-induced diseases (Supplementary Table S8). A few of them included genes with known functions reported in previous studies^3,20^. For example, the QTL located on the proximal end of chromosome 11 (Chr11:2 116 418 bp) associated with TMV resistance was associated with the *N* gene, a TIR-NBS-LRR protein introgressed through interspecific hybridization with *N. glutinosa^3,20^*. In addition, two QTLs, located on chromosome 7: 175, 557, 338 bp and chromosome 14: 2,809, 268 bp, associated with resistance to black shank (RBSH), were found to include *Sar8.2a* and *Sar8.2b* (Fig 3B), which were detected and fine-mapped previously^17^. Overall, these marker□trait associations represent a comprehensive catalogue of the allelic variations underlying differences in plant morphological, physiological, metabolic and disease-resistance traits, providing a deeper understanding of complex regulation and several candidates for future genetics research in this model plant species.

### Fine-mapping the causal variants underlying LW variation and functional characterization of the candidate gene Arf9

Motivated by broad interest in plant leaf development and its importance in light capture efficiency, we attempted to fine-map the QTL at Chr23: 148 211 202 bp associated with LW variation. A population of near-isogenic lines (NILs) was created by crossing two parents segregating at this QTL, Samsum (P1, T/T at Chr23: 148 211 202 bp) with wide leaves and K326 (P2, C/C at Chr23: 148 211 202 bp) with narrow leaves. Four generations of backcrossing were performed with K326 as the recurrent parent to create a BC4 population, and another four generations of self-pollination were performed to generate BC4F4 individuals. Based on the pattern of recombination and phenotype measurements for recombinant genotypes from BC4F2 and BC4F4 (Fig 5A, Supplementary Tables S13/S14), the QTL was fine-mapped to a 134 kb region between two markers, Chr23: 147 231 844 bp and Chr23: 147 366 149 (Fig 5A). There were only two SNPs between the two parents (Fig 5B) and one gene, NtZY23G02972, in this region. NtZY23G02972 is a homologue of *A. thaliana* auxin response factor 9 (*Arf9*), which widely exists in a number of crops but has not been functionally characterized. CRISPR□Cas9 knockout lines of NtZY23G02972 in *N. tabacum* var. *XHJ* showed an increase in LW by 6 cm (2.74×10^-3^; Fig 6 C-E, Supplementary Fig S7), suggesting that the nonfunctional allele of NtZY23G02972 was associated with wider leaves. Previously, the structure of ARF5 was characterized, and ARF DNA-binding domains are known to form a homodimer to generate cooperative DNA binding, which is critical for in vivo ARF5 function^21^. Here, we found that the second SNP on chromosome 23, 147 311 281 bp, located in the eighth exon of NtZY23G02972, altered the translated amino acid sequence from alanine (Ala_203_) to proline (Pro_203_), and the first SNP was located in the first exon of *Arf9*, altering the amino acid sequence from glutamic acid (Glu_29_) to glycine (Gly_29_). This second SNP was located inside the functional domain forming a homodimer (Fig 5F), while the first SNP was at the proximal end of the protein, which is less likely to have an impact on protein activity. Altogether, it is highly likely that a causal variant at chromosome 23, 147 311 281 bp, affected the formation of homodimers by changing the amino acid sequence from alanine (Ala_203_) to proline (Pro_203_) and increased LW. Our results revealed a novel gene regulating plant leaf development and provided a likely causal candidate for future functional genetic research.

**Fig 5.**
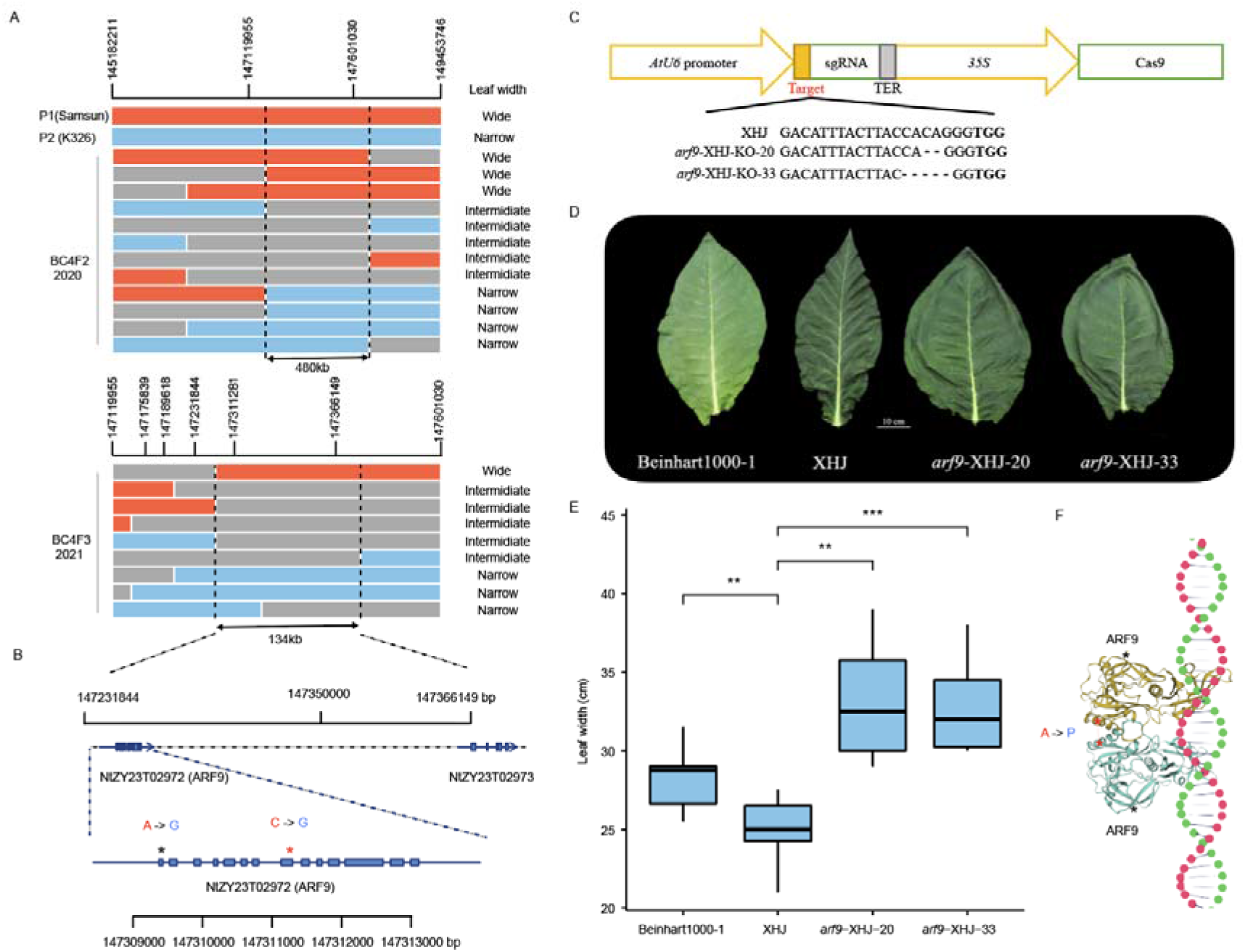
Fine-mapping and functional characterization of the QTL located at chromosome 23, 148 211 202 bp, associated with leaf width variation. **A).** Fine-mapping using near-isogenic lines (NILs). Each bar represents a unique genotype at this QTL region. Red represents the genotype of the donor parent (P1; Samsum) with wider leaves, while sky blue highlights the genotype of the recurrent parent (P2; K326) with narrower leaves. Unique offspring genotype classes from two generations, BC4F2 and BC4F3, are illustrated using coloured bars, and the line of origin is highlighted using red, sky blue, and grey for P1 homozygous, P2 homozygous, and heterozygous, respectively. **B).** Gene structure in the fine-mapped region and position of the two SNPs segregating in this population. A highly likely causal SNP is highlighted using red stars. **C).** Targeted sequences of NtZY23T02972 (*Arf9*) in CRISPR□Cas9 knockout lines. **D).** Illustration of the leaf width for a control line (Benhart 1000-1) and targeted line before (XHJ) and after (*arf9*-XHJ-20, *arf9*-XHJ-33) CRISPR□Cas9 knockout. **E).** Boxplot of the leaf width for a control line (Benhart 1000-1) and targeted line before (XHJ) and after (*arf9*-XHJ-20, *arf9*-XHJ-33) CRISPR□Cas9 knockout. **F).** A SNP altered the protein sequence of ARF from P > M, potentially affecting the formation of a homodimer to generate cooperative DNA binding.

**Fig 6.**
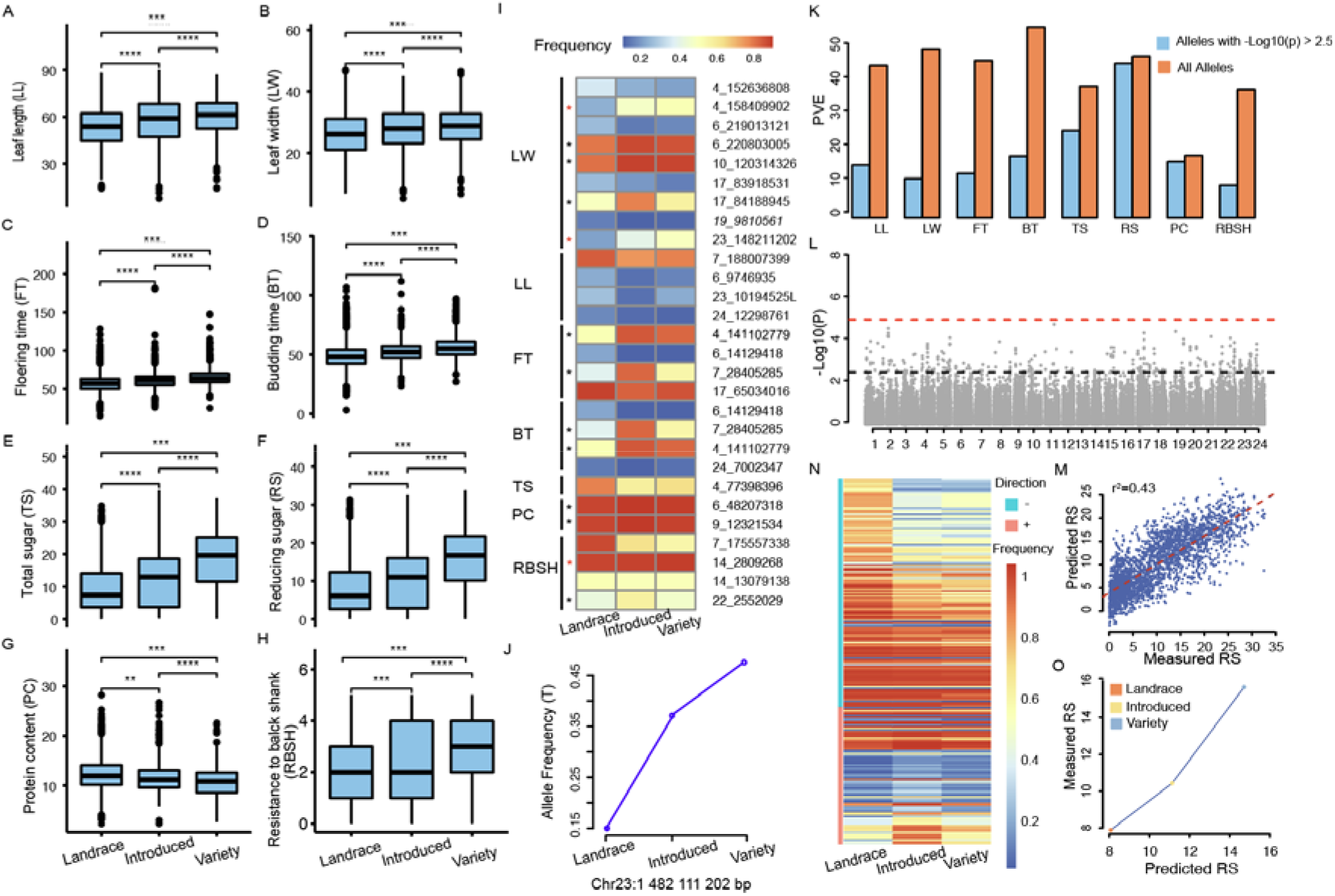
Signature of selection during selective breeding. **A-H)** Phenotype distribution of eight traits among different categories of materials (landraces, introduced varieties, and varieties) developed during the process of breeding. **I).** Frequency shifts of 28 QTLs associated with the variation in eight traits. **J).** Frequency shifts of the QTL associated with leaf width. **K).** The proportion of variance explained (PVE) by the alleles with −log10(P) values above 2.5 and all the alleles, estimated in a linear mixed model. **L).** Manhattan plot for RS association analysis. **M**). Scatterplot between predicted RS values based on 229 alleles with −log10(P) values above 2.5 and measured alleles. **N).** Frequency shifts of the 229 alleles with −log10(P) values above 2.5 in genome-wide association analysis of RS. **O).** Relationship between the mean of predicted RS level based on 229 alleles with −log10(P) values above 2.5 and measured RS values for three categories of plant materials developed during the process of breeding.

### Signature of positive selection and polygenic selection during the breeding process

When the germplasms were classified into landraces, introduced varieties (varieties developed from abroad) and Chinese local varieties (abbreviated as varieties hereafter) developed over a period of several hundred years, seven traits, LW, LL, FT, SL, RS, TS and RBSH, were found to have been continuously increased/decreased during the process of breeding (Fig 6 A-H). To test for evidence of selection, we evaluated whether the frequencies of favourable alleles associated with each trait increased over time. In total, 46.42% of the QTLs (highlighted using black/red stars in Fig 6I) showed an increase in allele frequency during the process of breeding. However, none of the alleles reached fixation (frequency >0.95), and only 17.00% of the QTLs showed allele frequencies greater than 0.8, indicating that there is great potential for future genetic improvement. Among these, 3 QTLs (10.71%) displayed lower allele frequencies in landraces than in introduced varieties, followed by varieties (highlighted using red stars in Fig 6I). For example, one QTL located at chromosome 23: 148, 211, 202 bp is associated with LW variation. Allele T at this locus increased LW by 1.11 cm (Fig 6J; P value = 1.67×10^-9^), and the frequency of the T allele continuously increased from 0.15 (landraces) to 0.38 (introduced varieties) and then to 0.47 (varieties; Fig 6J), suggesting that this allele is under positive selection.

In addition, polygenic scores were calculated using alleles with a −log10(p value) greater than 2.5, representing aggregated effects of minor-effect alleles. Overall, these minor-effect alleles (less than 0.5% of all SNPs) accounted for 23.12% to 95.56% (median = 33.40%, Fig 6K) of the kinship heritability, suggesting an important contribution of the polygenic background to the variation in these traits. Taking RS as an example, although no significant association was detected at the genome-wide significance threshold, aggregated effects of 229 (0.23%) alleles with −log10(p values) above 2.5 (Fig 6L) explained 43% of the phenotypic variation and 95.56% of the kinship heritability (Fig 6L/M). In total, 37.55% (86) of the alleles showed an increase in allele frequency from the landraces to the introduced varieties or varieties (Fig 6N). Although the magnitude of the frequency increases was small (mean = 0.07, Fig 6N), together they increased the level of reducing sugar from 8.07% to 14.71% from landraces to varieties (Fig 6O), demonstrating the power of polygenic adaptation in response to artificial selection. Overall, these results demonstrated the power of leveraging GenBank genomics for dissecting the genetic basis of complex trait variation and evolution during the process of selective breeding and provided a blueprint for future crop improvement.

## Discussion

*N. benthamiana*, an ancient allotetraploid species (~ 6 Mya) in the *Nicotiana* genus, experienced a significant reduction in chromosome number (24 >19) and genome size (~50%)^22^ after polyploidization. In contrast, over 90% of the parental genomic sequences are retained in the *N. tabacum* genome, and only 12 chromosome rearrangement events have been identified. This suggests that genome downsizing occurs on an evolutionary scale of several million years, and the stabilization of chromosome pairing during meiosis by chromosome rearrangement was likely to be the primary process in such a young polyploid (0.2 Mya). The high sequence similarity and high expression divergence between 1276 homologous genes (Supplementary Fig S3) indicate likely significant epigenetic remodelling for subgenome coordination.

In curing crops, such as tobacco and coffee, a primary target of breeding is balancing quality with certain morphological and phenological traits, whereby directional selection is relatively rare in comparison to that in field or horticulture crops. In addition, the history of selective breeding in tobacco is considerably short, being only ~ 200 years for flue-cured varieties and ~ 500 years for all other varieties. These factors likely explain the observed low divergence between major tobacco types, which in turn facilitated high statistical power for GWASs in *N. tabacum*. Future research with resequencing data is required to investigate the signatures of selection at a finer resolution.

We provided a comprehensive genotype-to-phenotype map in this model plant species and demonstrated the power of this roadmap in plant functional genetics by fine-mapping one of the detected QTLs. For example, LW is an important morphological trait associated with photosynthesis and is of great importance in light capture efficiency. By leveraging the genotype-to-phenotype map, we detected, fine-mapped and functionally validated a novel gene, *Arf9*, that regulates LW. Homologues of *Arf9* are widely present in a number of field crops, such as maize, rice, and wheat, and but not been functionally characterized. It is worthwhile to evaluate the function and potential of *Arf9* in major crop molecular genetic research and breeding applications.

In summary, we presented the first chromosome-level assembly of the *N. tabacum* allotetraploid genome and detected global genetic and phenotypic polymorphisms by sequencing and phenotyping an entire GenBank collection of 5196 *N. tabacum* germplasm resources. The extensive genotype and phenotype datasets, marker□trait associations, and candidate genes presented in this study will serve as a community resource for accelerating future comparative genomics, plant functional genomics and crop improvement research.

## Materials and Methods

### Plant materials, DNA extraction and sequencing

Total genomic DNA was collected and extracted from fresh leaves of *N. tabacum L*. var *ZY300* using the CTAB method. For PacBio long-read sequencing, 20 kb insertion SMARTbell libraries were constructed and sequenced on the PacBio Sequel II platform (Pacific Biosciences). For 10x Genomics sequencing, 1 ng of genomic DNA with a long sequence length (approximately 50 kb) was partitioned by a microfluidic chip on the Chromium platform, and 16-bp barcodes were introduced into droplets. For Illumina short-read sequencing, paired-end libraries with insert sizes of 350□bp were constructed and sequenced on the HiSeq PE150 platform. For Hi-C sequencing, DpnII-digested and cross-linked DNA was labelled with biotin and proximity-ligated to form chimeric junctions. Biotin-labelled samples were captured and sheared into 350-bp fragments. After terminal repair, A addition, joint connection, and library construction, Illumina sequencing was performed on the HiSeq PE150 platform. For Bionano sequencing, high-molecular weight DNA with a fragment distribution greater than 150 kb was isolated using Bionano sample preparation kits (Bionano Genomics). Genomic DNA was labelled using Direct Label Enzyme (DLE-1) and stained following BioNano protocols. Then, the labelled DNA was loaded into a nanochannel (Saphyr Chip, Bionano Genomics) and imaged using the Saphyr system (Bionano Genomics) following the Saphyr System User Guide.

Low-quality paired reads (reads with ≥ 10% unidentified nucleotides (N); > 10 nt aligned to the adapter, > 20% bases having a phred quality score <5 and putative PCR duplicates generated during the library construction process), which resulted mainly from base-calling duplicates and adapter contamination, were removed. In total, these steps yielded 1.79 Tb of high-quality data for chromosome-scale scaffold analysis.

### Genome assembly and scaffolding

For initial genome assembly, PacBio long sequence data were self-corrected to generate preassembled reads and assembled by the overlap-layout consensus algorithm^23^. The assembled contigs were further polished with Illumina short-read sequence data by the Pilon pipeline^24^. The polished contigs were further connected to generate super scaffolds using 10x Genomics linked-reads data using fragScaff software^25^. Scaffolds were anchored by linkage information, restriction enzyme site, and string graph formulation with the package LACHESIS^26^. Hi-C data were used to further improve the quality of the assemblies with HiC-Pro software (v.2.10.0). The placement and orientation errors exhibiting obvious discrete chromatin interaction patterns were adjusted manually. A final assembly was generated with Bionano Solve (v3.5.1). Chromosomes were named by mapping SSR markers from a widely used linkage map^27^ to unify the physical map and linkage map.

### Repeat and gene annotation

Extensive de novo TE Annotator (EDTA v1.9.3)^28^, a pipeline combining homology-based and de novo prediction methods using LTRharvest^29^, LTR FINDER^29^, LTR retriever^30^, TIR-learner^31^, Helitron Scanner^32^, and Repeat Modeler^33^, was used to annotate TEs, estimate TE insertion time and generate a species-specific library for gene annotation. Structural annotation of genes was conducted through a combination of homology-based, transcriptome-based and ab initio-based methods. For homologue prediction, protein sequences of plants, including *N. tabacum* (https://solgenomics.net/ftp/genomes/Nicotiana_tabacum/edwards_et_al_2017/annotation/), *Solanum tuberosum* (GCF_000226075.1), *Solanum lycopersicum* (GCF_000188115.4), *Coffea canephora* (PRJEB4211_v1), *Gossypium hirsutum* (GCF_000987745.1), *A. thaliana* (GCA_000001735.1) and *Vitis vinifera* (http://jul2018-plants.ensembl.org/Vitis_vinifera/Info/Index), were used as queries to search against the ZY300 genome using GeneWise (v.2.4.1)^34^. For transcriptome-based gene prediction, trimmed RNA-sequencing reads from stems, roots, leaves, anthers, flowers, and axillary buds were de novo assembled using Trinity (v2.1.1) ^35^. To further improve the annotation, RNA-Seq reads from different tissues were aligned to the ZY300 genome using TopHat (v2.0.11) with default parameters to identify exons and splice junctions. The alignment was used as input for Cufflinks (v2.2.1) ^36^ for transcript assembly with default parameters. For ab initio prediction, we used Augustus^37^, GlimmerHMM^38^, SNAP^39^, GeneID (v1.4)^40^ and Genscan^41^ to predict gene structure. Finally, all predictions of gene models generated from these approaches were integrated into a consensus gene set using EVidenceModeler (v1.1.170)^42^. After prediction, PASA^43^ was used to update isoforms to gene models and to produce a final gff3 file with three rounds of iteration.

For functional annotation, predicted protein-coding genes were aligned to multiple public databases, including NR, Swiss-Prot, TrEMBL75, COG, and KOG, using NCBI BLAST□ + □v.2.2.31^44^. Motifs and domains were annotated using InterProScan (release 5.32-71.0)^45^. Gene Ontology (GO) terms and KEGG pathways of predicted sequences were assigned by InterProScan and KEGG Automatic Annotation Server^45,46^, respectively.

### Subgenome partitioning and synteny analysis

Illumina short-read sequences from *N. sylvestris* (accession ID: ERR274529) and *N. tomentosiformis* (accession ID: ERR274543) were downloaded from the NCBI SRA database and aligned to the ZY300 reference genome using bwa-mem^47^. SAMtools^48^ was used to count the number of reads mapped to the ZY300 genome from each of the two parental genomes, and the average read depth in 1-Mb windows was calculated using a customized R script. First, a preliminary parental origin assignment was made using the window-averaged read ratio. Second, for windows containing breakpoints, manual curation was performed in Integrative Genomics Viewer^49^ (IGV) to pinpoint the exact location. WGDI^50^ was used to detect synteny blocks based on collinear genes, and the R package circlize^51^ was used to illustrate blocks between subgenomes after merging large synteny blocks.

### Subgenome expression bias analysis

Four hundred RNA sequencing datasets were randomly selected from the NCBI SRA database (accession ID available in Supplementary Table S16) to quantify the level of expression bias for homologous genes from two parental genomes. Raw data were downloaded and trimmed using fastp^52^ with the default parameters and subsequently aligned to the ZY300 reference genome using HISAT2^53^. The expression level was quantified as FPKM using Stringtie^54^. Fold change between homologous genes was calculated as the ratio between averaged FPKM values across the 400 samples.

### Genotyping by sequencing, read mapping, and variant calling

DNA extraction was performed using CTAB methods. After quality control using Nanodrop and Qubit, DNA samples were digested with *NlaIII* + *MseI*, barcoded and purified using AMPure XP beads. Quality-controlled libraries were pooled and sequenced on the Illumina NovaSeq platform. Raw reads were trimmed using fastp^52^ following the default parameters and subsequently aligned to the ZY300 reference genome using bwa^47^. Samtools^48^ mpileup was used for variant calling. The called SNPs were initially filtered for read depth, call rate, and minor allele frequency with vcftools^55^ on the basis of average read DP > 2 and MAF > 0.03, individual missing rate < 0.5, and site missing rate <0.5. The filtered genotypes were then imputed using beagle^56^. The imputed SNPs were further filtered for a MAF greater than 0.03, and 95,308 SNPs remained for downstream analysis.

### Population structure analysis

To further assess the relatedness between individuals, principal component analysis was performed using the identity-by-state (IBS) genetic distance matrix calculated in PLINK (v1.90)^57^, and principal component analysis was performed using the *princomp* function in the R base package. After grouping samples, VCFtools (v0.1.16)^55^ was used to calculate the population differentiation coefficient (Fst) among major tobacco types and genetic clusters.

### Genome-wide association analysis

Genome-wide association (GWA) analysis was conducted for 43 traits (phenotype measurements are available in Supplementary Table S17) using a linear mixed model implemented in the *mlma* module of GCTA^58^. A subsequent conditional analysis implemented in the *cojo* module of GCTA^58^ was performed to screen for independent association signals. As the LD was extensive in this population, assuming that all tested markers were statistically independent and deriving a Bonferroni-corrected significance threshold would have been too conservative. Therefore, we estimated the effective number of independent markers (Me)^59^ and derived a less conservative threshold following 0.05/Me (1. x 10^-5^ equivalent to −log_10_ (*p* value) = 5).

### Polygenic selection analysis and genomic prediction

We predicted the aggregated effects of all the SNPs with a −log10 (p value) above 2.5 using the following model.

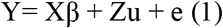

where Y is a column vector of length n containing phenotype measurements. X is a matrix of n rows and 1 column on 1s, representing the population mean. u is a random effect vector of the polygenic effects (polygenic scores) representing the aggregated effects of all the SNPs for the n individuals. Z is the corresponding design matrix obtained from Cholesky decomposition of the kinship matrix G, estimated on the basis of the markers with a −log10 (p value) above 2.5 using the *A.mat* function in the R package rrBLUP ^60^. The Z matrix satisfies ZZ’=G; therefore, 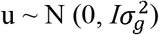. e is the residual variance with 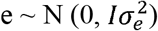. The proportion of variance explained by these SNPs was estimated as 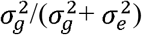. Model fit was assessed in the R package rrBLUP ^60^.

### Fine mapping and experimental validation of candidate genes

A population of NILs was developed by marker-assisted backcrossing using Samsun (P1, T/T at Chr23: 148 211 202 bp) as the wide-leaved donor parent and K326 (P2, C/C at Chr23: 148 211 202 bp) as the narrow-leaved recurrent parent. First, a total of 1694 BC_4_F_2_ individuals from the K326×Samsun population were genotyped using five markers from Chr23: 145 182 211 bp and Chr23:149 453 746 bp (Supplementary Table 13), and then selected recombinants were self-pollinated to create BC_4_F_3_ lines for further evaluation. This narrowed the QTL to a region between Chr23: 147 119 955 and Chr23: 147 360 000bp. An additional 3017 BC_4_F_3_ individuals derived from the population were further screened using seven SNP markers in this narrow region (Supplementary Table 13), and the BC_4_F_4_ lines of recombinants were phenotyped for further fine mapping. Two sgRNAs (small guide RNAs, Fig 5C) were designed to target the candidate gene NtZY23T02972 based on the assembled ZY300 genome. The vectors were constructed and transformed into the receptor XHJ. The genotype of gene-edited lines was identified by PCR amplification and Sanger sequencing. For fine mapping, LW was recorded by measuring at least 15 plants per line during flowering in Sanya (E 108.56, N 18.09), China. For transgenic experiments, phenotypes of knockout lines and the wild type were investigated in Sanya (E 108.56, N 18.09), China. Each experiment had two replicates with at least five independent plants per replicate. The mean LW of at least five plants per replicate was used for further analyses.

## Author Contributions

A.Y., L.C. and Y.C. designed and supervised this study. Y.Z., S.C., M.R., G.L., and Y.L. collected the data and implemented the analysis. H.S. contributed to the statistical analysis. Y.L., X.Z., Y.T., C.J., Z.L., D.L., Y.S., Z.X., and R.G. conducted the field experiments and phenotype measurements. Q.F. and Y.W. collected the germplasms. G.Y. and L.F. offered valuable input during the design and implementation of this study. Y.Z., S.C. and L.C. wrote the manuscript.

All authors have read and approved the manuscript and have no conflicts of interest to declare.

## Acknowledgements

We thank Yan Ji for her efforts in improving the figures and tables. We thank Wei Zhao and Ying Guo for their constructive comments to improve the manuscript. We thank Shilin Tian for his valuable input during the design and implementation of this study.

## Data availability

All sequence data will be available from the NCBI upon acceptance of this manuscript.

All phenotype measurements and metadata of samples are included in supplementary tables.

